# Understanding the survival of Zika virus in a vector interconnected sexual contact network

**DOI:** 10.1101/518613

**Authors:** Tanvir Ferdousi, Lee W. Cohnstaedt, D. S. McVey, Caterina M. Scoglio

## Abstract

The recent outbreaks of the insect-vectored Zika virus have demonstrated its potential to be sexually transmitted, which complicates modeling and our understanding of disease dynamics. Autochthonous outbreaks in the US mainland may be a consequence of both modes of transmission, which affect the outbreak size, duration, and virus persistence. We propose a novel individual-based interconnected network model that incorporates both insect-vectored and sexual transmission of this pathogen. This model interconnects a homogeneous mosquito vector population with a heterogeneous human host contact network. The model incorporates the seasonal variation of mosquito abundance and characterizes host dynamics based on age group and gender in order to produce realistic projections. We use a sexual contact network which is generated on the basis of real world sexual behavior data. Our findings suggest that for a high relative transmissibility of asymptomatic hosts, Zika virus shows a high probability of sustaining in the human population for up to 3 months without the presence of mosquito vectors. Zika outbreaks are strongly affected by the large proportion of asymptomatic individuals and their relative transmissibility. The outbreak size is also affected by the time of the year when the pathogen is introduced. Although sexual transmission has a relatively low contribution in determining the epidemic size, it plays a role in sustaining the epidemic and creating potential endemic scenarios.

## Introduction

Zika virus (ZIKV) is predominantly vector-borne and the outbreaks are strongly dependent on the mosquito vector density which is influenced by the climatic conditions^1^. There have been substantial evidence2 and reports3 of sexual transmission of ZIKV. Although ZIKV epidemic models mostly focus on the mosquito-vectored transmission, multiple studies have quantified the contribution of sexual transmission^4–8^. Modeling analysis has helped to determine the relative importance of different transmission mechanisms. To enhance our understanding on how the combination of both vectored and sexual transmission can help sustain the pathogen during the vector free seasons (e.g. winter), we develop a novel epidemic model that interconnects a homogeneous mosquito vector population with a heterogeneous sexual contact network.

Zika virus is a positive-sense single-stranded RNA virus of the Flaviviridae family^9^. It is related to other flaviviruses such as dengue, yellow fever, West Nile and Japanese encephalitis viruses. ZIKV infection symptoms include acute fever, maculopapular rash, arthralgia and conjunctivitis. However, only about 20% of infected people suffer any significant symptoms^10^. A large proportion of asymptomatic cases poses a major problem in determining the outbreaks to initiate early response measures. ZIKV infection in pregnant women can result into congenital microcephaly, a severe birth defect^11^. The transmission of ZIKV occurs primarily by infected mosquito vectors such as *Aedes aegypti* and *Aedes albopictus.* However, it can also be spread via infected semen and blood^12^. The first autochthonous case of Zika in the mainland US was reported in July 2016, in Miami, Florida. As of April 2018, 5, 676 cases have been reported in the US States^13^. In 2016, 224 autochthonous cases have been reported where the state of Florida itself is responsible for 97% of those cases, and the remaining 3% cases occurred in Texas^13^. Due to the complex nature and multiple pathways of pathogen transmission, several epidemic parameters are still unknown.

There have been numerous attempts in modeling the outbreaks of ZIKV. Several models only considered vectored transmission^1,14,15^ and several others considered both sexual and vectored transmission routes^4,5^. Vertical transmission has also been modeled in a work of Agusto et al.^16^. Olawoyin et al. analyzed the effects of multiple transmission pathways and concluded that, secondary transmision mechanisms increase the basic reproductive number and cause the outbreaks to occur sooner compared to the vector-only transmission^17^. The work of Gao et al. concluded that sexual transmission contributes about 3.044% of the overall transmission^4^. Moghadas et al. also found similar contribution from sexual transmission and they emphasized the importance of asymptomatic transmission^18^. Maxian et al. concluded that this contribution is too minor to sustain an outbreak, as they found that sexual transmission contributes about 4.8% to the basic reproductive number, *R*_0_^6^.

They also concluded that sexual transmission indirectly helps the vectored transmission by increasing the pool of infectious individuals. The relatively higher prevalence of infection in the female population compared to male population was analyzed by Pugliese et al.^19^, who concluded that higher susceptibility of females compared to their male counterparts in the case of sexual transmission could be the reason. The work of Sasmal et al. estimated that sexual transmission contribution could be as high as 15.36% for sexual risk-stratified population^20^. Another study concludes that existing research could potentially underestimate the risk of sustained sexual transmission by an order of magnitude^21^. It has been found that Zika virus can sustain in semen for a long time after it disappears from blood^22^. Hence, even though it is clear that sexual transmission alone can’t sustain an outbreak, we don’t have clear information about the persistence the pathogen in a human host network without the help of vectors. Limited information on the reduction of transmissibility for both asymptomatically infected hosts and during the extended sexual transmission period adds to the complexity in understanding how the outcomes are affected.

In this paper, we develop an individual-based (i.e., node-based) network model, which is different from basic compartmental models as it features heterogeneous mixing. Each individual (i.e., a node) is connected to a specific number of individuals and this number can be very different. In the context of real world sexual networks, this is realistic. Using survey data23 on sexual behavior, we develop a network generator tool that can produce random networks with properties that closely matches the given data. As the vector population is much larger than the host population and vectors don’t transmit between them, it is unnecessary to use node-based models for vectors. We employ a homogeneous population model for the vectors. We interconnect these two populations (hosts and vectors) via the use of infection rate parameters where the infection of one population depends on the number of infected in the other population. There has been several works on epidemic spreading in interconnected networks^24^. However, due to the multi-host (one being the vector) scenario, our model is built differently from conventional interconnected networks. Our goal is to incorporate heterogeneity in the host population but reduce unnecessary computational overhead by interconnecting with a homogeneous vector population. Our model also differentiates between symptomatically and asymptomatically affected individuals by assigning them different states. We furthermore, incorporate the extended sexual transmissibility in our model. In the host network sub-model, each host node can be in either one of the seven states: *SEI_s_I_a_J_s_J_a_R* (Susceptible, Exposed, Infectious (Symptomatic), Infectious (Asymptomatic), Convalescent (Symptomatic), Convalescent (Asymptomatic), and Recovered). The host population is also divided based on gender, sexual orientation, and age. In the vector population sub-model, the population is modeled into three compartments: *S_V_E_V_I_V_* (Susceptible, Exposed and Infected). We only use birth-death demography for the vector model and the climatic variation is incorporated into the vector birth rate, which is one of the variable parameters. The simulation model is based on Gillespie’s Stochastic Simulation Algorithm (SSA^)25^. However, we have a few variable rate parameters in our model, which prompted us to use the non-Markovian Gillespie Algorithm (n-MGA)26 to simulate our model, which we implement using a modified version of the GEMF27 tool.

The contributions of this paper are threefold: i) we propose a novel interconnected model to evaluate host-to-host and host-to-vector-to-host pathogen transmission, ii) we develop a sexual contact network generator based on aggregate data, and iii) using our implementation of the non-Markovian Gillespie Algorithm (n-MGA) we examine survival of pathogen in the host network. We also perform sensitivity analyses on key model parameters to evaluate their role in disease outcomes. The interconnected model is presented in the model formulation section, three independent approaches in evaluating model outputs are presented in the results section, and the methods section contains the details on the simulation model.

## Model Formulation

### Interconnected Population Model

We propose an interconnected population model to investigate the spread of ZIKV within the human host and the mosquito vector population during an outbreak. A basic diagram of the model is shown in Figure 1. It has two major components: the vector population marked by the cloud on top, and the host contact network marked by the solid circle nodes and the solid edges. The vector population is assumed to be in the vicinity of the host population such that each mosquito vector can bite any of the hosts (indicated by dashed lines).

**Figure 1.**
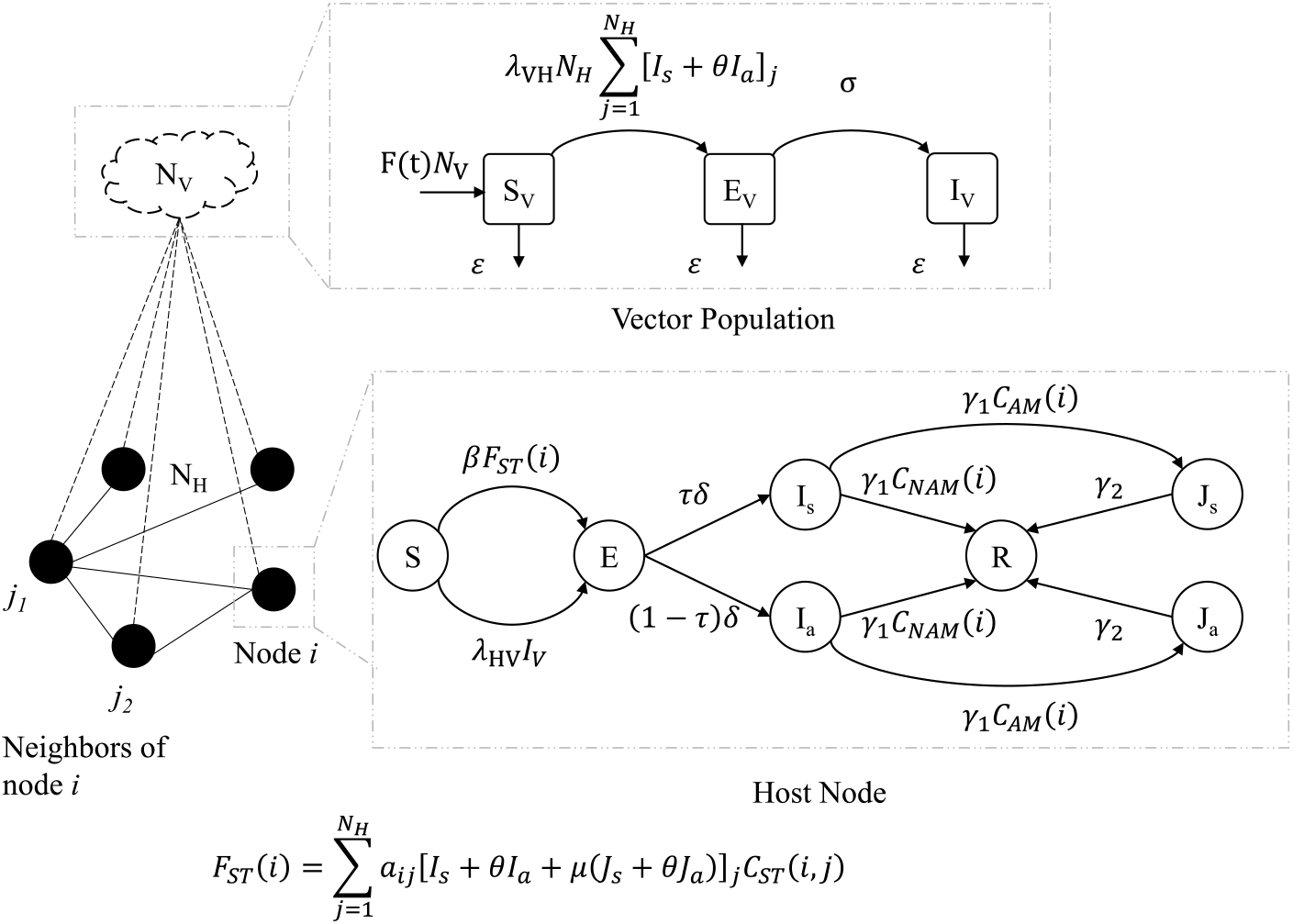
Coupled population model for ZIKV. The black solid circles indicate host nodes (individuals) and the cloud shape above indicates the vector population. The solid edges connecting the nodes indicate the sexual contact network and the dashed edges indicate contacts of the vector population with the host nodes. A host node can be in any of the seven states (*SEI_s_I_a_J_s_J_a_R*) and the entire vector population is divided into three compartments (*S_V_E_V_I_V_*).

Each host node can be in either one of the seven states: Susceptible (*S*), Exposed (*E*), Infectious (Symptomatic) (*I_s_*), Infectious (Asymptomatic) (*I_a_*), Convalescent (Symptomatic) (*J_s_*), Convalescent (Asymptomatic) (*J_a_*) and, Recovered (*R*). The convalescent states were used to model the extended presence of ZIKV in semen as reported by previous works^22^. People in *J_s_* or *J_a_* states cannot transmit infections via vectors, they can only transmit sexually. The vector population is divided into three compartments: Susceptible (*S_V_*), Exposed (*E_V_*) and, Infected (*I_V_*). The unidirectional arrows indicate transition from one state/compartment to another. The symbol adjacent to each arrow indicates the rate of that particular transition. The set of all these symbols constitute the model parameters, and they are listed in Table 1. This table also lists the nominal values and the range of values to be used for the simulation experiments. Some of these values are not readily available in the literature and we performed calibrations to determine acceptable ranges of values to conduct the experiments. We also performed sensitivity analyses with several critical parameters to evaluate model behavior. We have also used boolean condition functions marked by the symbol, *C_x_*(*n*) as shown in Figure 1. The subscript ‘*x*’ indicates which condition a node *n* has to satisfy. If a condition is met, the condition function returns True (= 1), otherwise it returns False (= 0). For example, *C_AM_*(*i*) will return 1 if node *i* is an adult male (AM), it will return 0 otherwise. The conditions that we have used are: *AM* = Adult Male, and *NAM* = Not Adult Male. It should be obvious that for any *i, C_AM_*(*i*) is the complement of *C_NAM_*(*i*). *F_ST_* (*i*) is the multiplier component function of the force of infection for sexual transmission from other nodes to node *i*. The symbols *I_s_, I_a_, J_s_* and, *J_a_* in the formulation of *F_ST_* (*i*) are host state indicators for neighbors (*j*) of node *i. C_ST_* (*i, j*) is the sexual transmission condition function. It only returns True (1) if *i* is an adult and *j* is an adult male. Here, *j* is considered the transmitter and *i* is consider the recipient of infection.

**Table 1.**
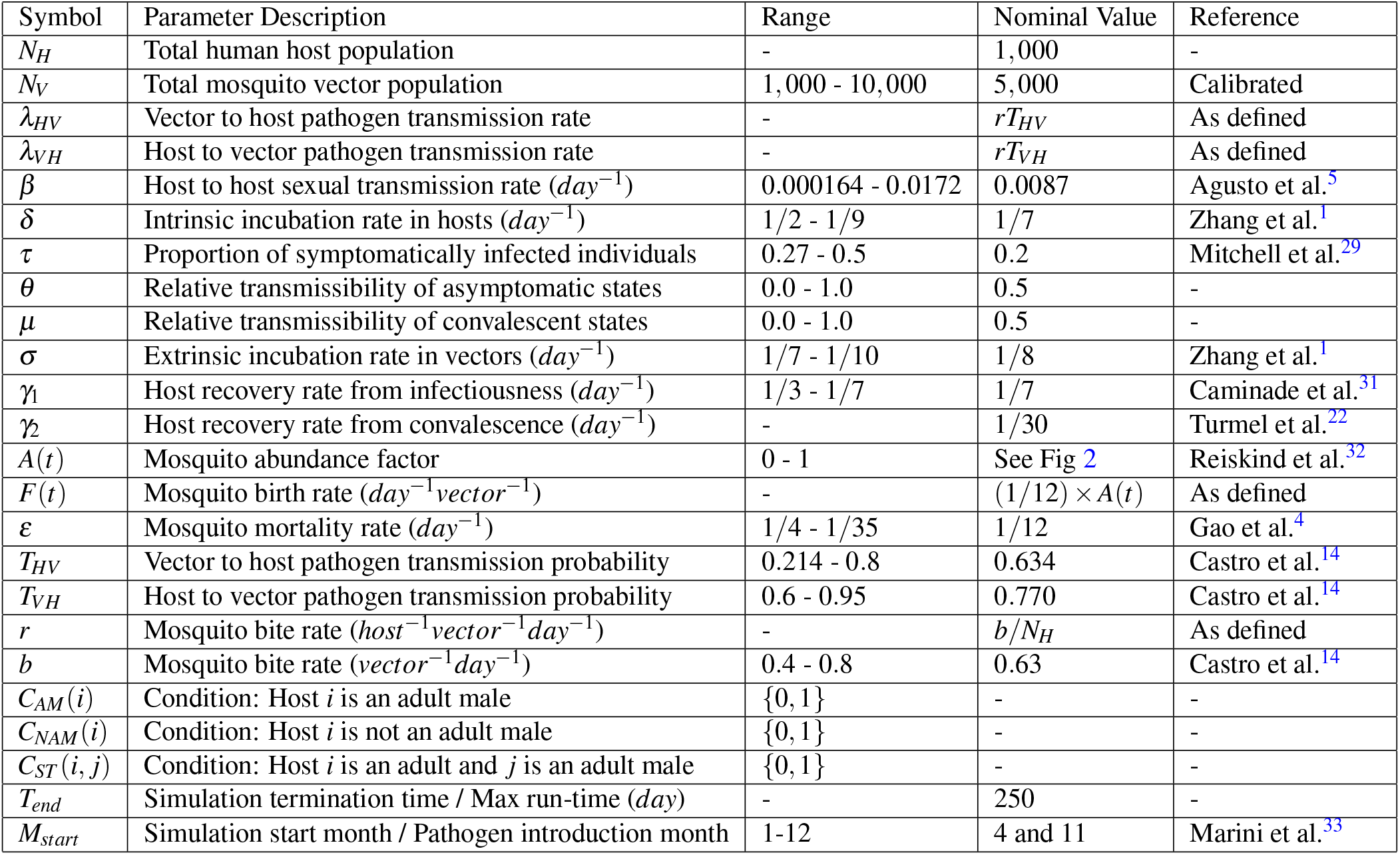
Epidemic model parameters and functions.

As indicated in Figure 1, a susceptible host node can get infected (*S* → *E*) via two mechanisms: i) getting bitten by an infected mosquito (at the rate *λ_HV_*) or ii) having sexual contact with an infected or convalescent (*I_s_, I_a_, J_s_* or, *J_a_*) host (at the rate *β*). We use the adjacency matrix representation of a graph, where 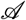 is the adjacency matrix. An element 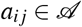 indicates the connection/link between node *i* and node *j.* If a link exists, then *a_ij_* = 1, otherwise *a_ij_* = 0. The sexual transmission rate, *β* was computed using the formula *β* = 1 - (1 - *α*)^*n*^, which was previously used in the work of Agusto^5^. Here, *α* is the probability of transmission per coital act with an infected partner. As data specific to Zika virus is not available, we assume a range of *α* between 0.001 and 0.1^28^. The exponent *n* is the average number of coital acts per unit time. Some estimated data indicate that people have sex an average of 60 times per year^23^. Hence, we estimate *n* = 60/365 = 0.1644 per day. The range of *β* for the range *α* is 0.000164 to 0.0172. Once a human host is infected, he/she can either develop symptoms or not show any sign of infection at all. Hence, there are two possible transitions from the exposed (*E*) state: i) a proportion, *τ* of the people becomes symptomatically infected (*I_s_*) and ii) a proportion, 1 – *τ* of the people becomes asymptomatically infected (*I_a_*). The proportion of symptomatic ZIKV infections is claimed by previous works10 to be about 20%. However, one recent work^29^ claims that this proportion could be higher ranging from 27% – 50%. For our analysis, we take this into consideration as shown in Table 1. Once someone is infected, they will stay infected for an average duration of 1/*γ*_1_. However, once the virus disappears from blood, the adult male hosts will go through some extended period of infectiousness which is modeled by the convalescent states (*J_s_* and *J_a_*). An adult male in the convalescent state may only transmit sexually and would recover after an average duration of 1/*γ*_2_. The infectiousness of the 4 states (*I_s_, I_a_, J_s_* and, *J_a_*) are not considered the same. The relative transmissibility of the asymptomatic states is represented by *θ*. The relative transmissibility for the convalescent states is denoted by *μ*. Although, symptomatically infected people could be spreading less if they are seeking medical attention, we do not reduce the transmissibility for the symptomatic cases in this model. Historically, it has been seen that, possible negligence associated with ZIKV has contributed to sexual transmissions^30^. A healthy mosquito vector can get infected with some nonzero probability only if it bites an infectious human (*I_s_* or *I_a_*) (at the rate *λ_VH_*). The time dependent per-capita mosquito recruitment rate is represented by *F*(*t*) where, *F*(*t*) = (1/12)*A*(*t*). Here, 1/12 is a fixed birth rate assumed to be the same as the mortality rate (*ε*). The mosquito abundance factor *A*(*t*) is the seasonality parameter which incorporates seasonal variations into the model.

To generate a realistic sexual network, we develop a network generator tool based on the ‘configuration model’^34^. Our tool takes as input data of sexual behavior, age, gender, and produces representative random networks. Details are featured in the Methods section. An example network generated by our tool is given in Figure 3. In our model, we also intend to incorporate the temporal variations of the climate in different seasons and the consequences of such variation on the mosquito vector population. To simplify this task, we use a vector abundance factor (*A*(*t*)) expressed as a fraction in the range [0,1]. This parameter was derived from data points used in the work of Monaghan et al.^35^, which they originally extracted from the works of Reiskind et al.^32^. For our simulations, we use vector abundance data from Miami, Florida. The mosquito abundance factor over the course of 12 months is plotted in Figure 2. The data originally had 12 sample points (monthly values). As the model requires the abundance factor on a daily basis, we use linear interpolations between data points of adjacent months.

**Figure 2.**
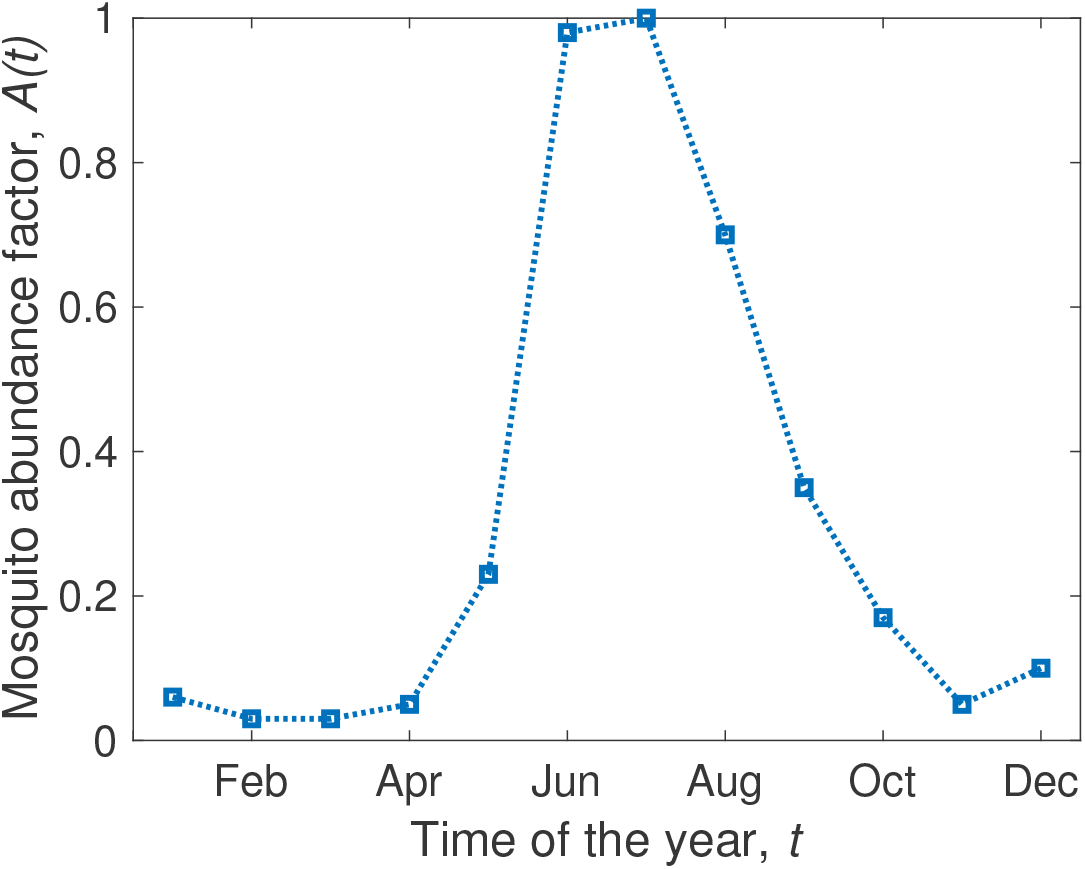
Variation in mosquito vector abundance with time. Due to changing temperature, rainfall and other climatic factors, mosquito abundance varies throughout the year. The squares show the data points and the dotted lines are linear interpolations. The data represent Palm Beach, Florida where the vector abundance peaks during June-July. The plot was constructed from data observed in Palm Beach, FL over 27 four-week periods from 2006-2008^32,35^.

**Figure 3.**
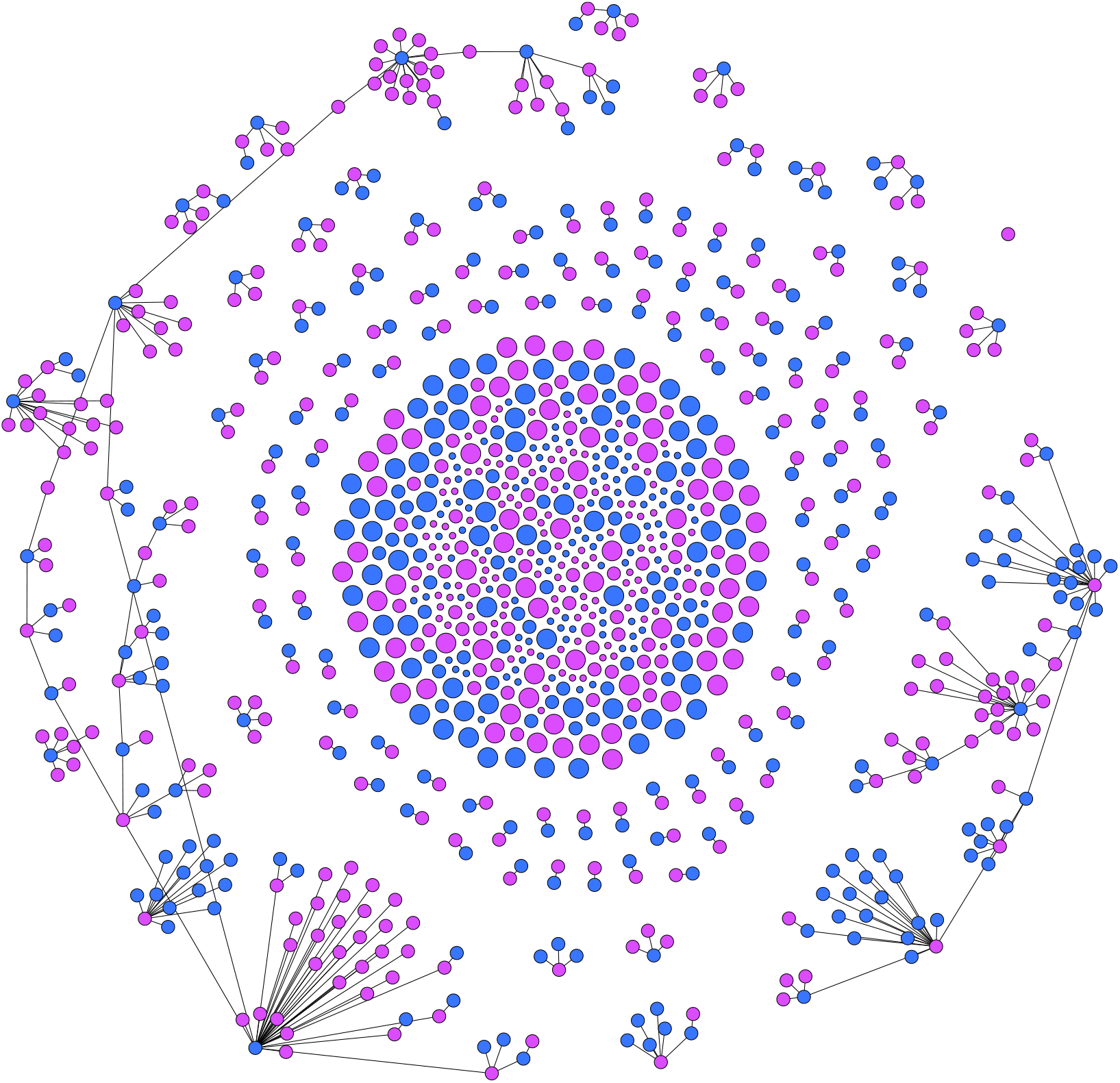
Generated sexual contact network based on sexual behavior^23^. The given network has *N* = 1000 nodes. The Methods section contain the details on how it is generated. The nodes are presented in two distinct colors and three distinct sizes. The males are marked as blue and the females are marked as pink. The three node sizes correspond to Children (small), Adults (medium) and Elderly (large) people. This particular instance of the network has an average node degree of 0.803 whereas the 64% adult population have an average degree of 1.26. This particular instance of network has 463 male and 537 female host nodes. The edge density of this network is 0.0008038. There are 600 connected components and the largest one consists of 117 nodes.

### Simulation Tool

If the transition rate parameters are constant, the overall stochastic model is a Markov process with Poisson arrivals and exponential inter-event times. A stochastic algorithm such as the Gillespie SSA25 can be used to perform simulations in most of these cases. The well-developed GEMFsim27 can be used to solve such stochastic spreading processes in the networks. However, our model requires a few changes before such simulations can be performed.

In our model, the transition rate parameters are not constant. The mosquito birth rate is a seasonally dependent parameter that varies according to the given vector abundance input data. The vector population compartments are changing with time during the outbreak simulation, which change the host infection rate *λ_HV_I_V_*. Similarly, the host infected populations are also changing with time which modify the vector infection rate 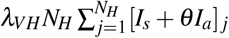. These constitute a set of exogenously varying parameters (i.e., varying due to external forces/catalysts). As a consequence, the processes are no longer Poisson, rather these are called non-homogeneous Poisson. To cope with this issue, we use the non-Markovian generalized Gillespie Algorithm (nMGA)26 which assumes general renewal processes (which is more generic) that allows exogenous variations of parameters. We modified the existing GEMFsim27 accordingly, and added a vector compartmental model solver on top of it. The vector compartmental solver is effectively coupled to the modified network-based nMGA solver via the infection rate parameters (*λ_HV_I_V_* and 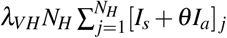), where a parameter in one population depends on some quantity in the other population.

## Results

The simulation tool uses the parameters listed in Table 1, the initial conditions, the seasonal variation data, the population data, and the contact network representation. The initial condition is a single infected vector (*I_V_* (0) = 1) while keeping the remaining vector population and the entire host population susceptible (*S*). We specify the pathogen introduction month (*M_start_*) and the max run time (*T_end_*). For example, if we set *M_start_* = 5 and *T_end_* = 100, the simulation will start from 1^st^ of May and run up to the 2^nd^ week of August. A simulation can terminate before *T_end_* if the outbreak dies out. As we are running stochastic simulations, we smooth out our results by averaging over 500 repetitions. The population data contains the distribution of genders, sexual orientations and age groups, used to generate the contact network. For the survival analysis, we introduce the pathogen in November (*M_start_* = 11) to indicate the start of winter. For the sensitivity analysis, we introduce the pathogen into the population in April (*M_start_* = 4), which is the most likely scenario for the 2016 Miami-Dade outbreak^33^.

### Seasonal Analysis

First, we explore the effect of seasonal variations on the epidemic progression. We use the nominal parameter values defined in Table 1 and run the simulations for three distinct pathogen introduction months, *M_start_*. Taking some cues from the seasonal patterns of Miami, FL shown in the Figure 2, we choose to run independent scenarios starting on 1^st^ of April (*M_start_* = 4), 1^st^ of August (*M_start_* = 8), and 1^st^ of October (*M_start_* = 10). The averaged results of 500 simulations are shown in Figure 4.

Without any intervention, we observe a large outbreak (more than 50% of the population affected) if pathogen is introduced on 1^st^ of August. For the remaining two cases, the outbreak sizes are comparatively much smaller. This can be explained using the seasonal variations of Figure 2. In Miami, the mosquito abundance is high during the beginning of August. This contributes to a boost in the infected vector population compared to the other two dates.

We also observe some interesting behaviors in the outbreak dynamics. Despite the vector abundance having a large positive slope during the month of April and a large negative slope during the month of August, Miami suffers a larger outbreak in the August introduction compared to the April introduction. Figure 4b shows that the infected vectors die out soon in the April outbreak although the upcoming summer (positive slope in abundance) sustains healthy vectors for a long time. In the case of the August introduction, the infected vectors rise rapidly during the first 50 days (Figure 4d). Although the vector population declines rapidly after 50 days due to unsuitable climates of the upcoming winter, the initial surge in infected mosquitoes causes a large outbreak. The results indicate that *the suitability of the climate during pathogen introduction plays a major role in determining the size of the outbreak.* The impacts of climatic changes in the following months are minimal.

**Figure 4.**
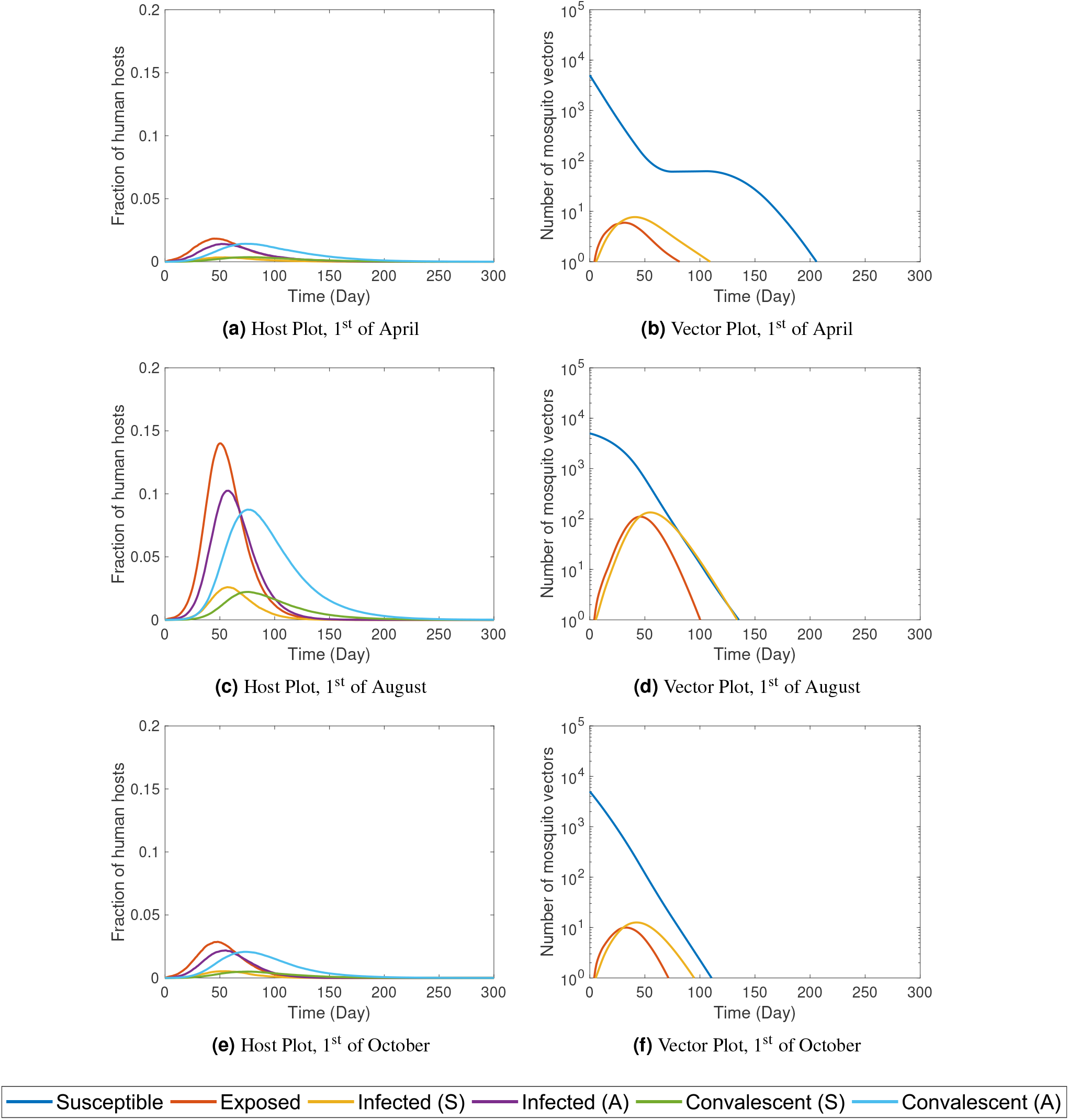
The time series plots of a ZIKV outbreaks for three different pathogen introduction months, *M_start_* in Miami, FL. The left column contains the host plots and the right column contains the corresponding vector plots. The three rows indicate three different pathogen introduction times: 1^st^ of April, 1^st^ of August, and 1^st^ of October respectively. The hosts in different states are expressed as fractions of the total population. The number of vectors in different compartments are expressed in log scaled axes. Here, we present 5 out of the 7 host states (*E, I_s_, I_a_, J_s_* and, *J_a_*) and the all three vector compartments (*S_V_, E_V_* and, *I_V_*) which are marked with distinct colors described by the legends at the bottom. All plots are averages of 500 independent stochastic simulations.

### Survival Analysis

As a part of the survival analysis we will be observing several measured quantities: the epidemic/infection lengths, the pathogen survival period, and the epidemic attack rates. Here, we define them first before discussing the results.

#### Host Infection Length (T_HL_)

The host infection length is defined as,

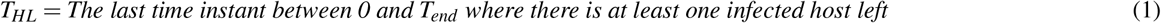

It is the last day on the outbreak time-line an infected host can be found. For this property, the infected hosts who are asymptomatic or in the convalescent state are also considered infected. In this article, when we use the term *“epidemic length”,* we imply host infection length *T_HL_* unless otherwise mentioned.

#### Vector Infection Length (T_VL_)

The vector infection length is defined as,

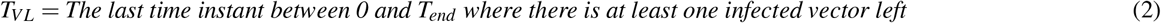

It is the last day on the outbreak time-line an infected vector can be found. In typical outbreak situations, infected vectors die out before all the infected hosts recover.

#### Pathogen Survival Period, (T_PS_)

The vector free pathogen survival period is defined as,

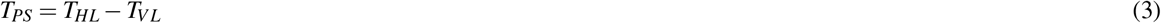

It measures how long pathogen can survive in the host population without the presence of vectors.

#### Epidemic Attack Rate (AR)

The epidemic attack rate (*AR*) is defined as,

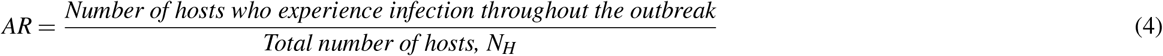

The value of *AR* is in the range [0, 1]. We sometimes express this quantity as a percentage instead of a fraction. For example, an *AR* of 0.3 means 30% of the population were infected during the outbreak and the remaining 70% never experienced any infection.

To see how some of the above mentioned quantities relate to sexual transmission, we vary the sexual transmission rate, *β* in the range mentioned in Table 1. We find that varying *β* does not noticeably increase or decrease the epidemic length (Figure 5a), which remains between 2-4 days for the range of *β* in consideration. Outbreaks in most of these cases last for about 158 days. The effect of increased sexual transmission in modifying pathogen survival is also minor (Figure 5b), being within 74-78 days with an average of 76.75 days. A similar situation arises for the attack rate as well (Figure 5c), with the mean attack rate being 15.12%. If there is no sexual transmission (*β* = 0), the attack rate is 14.48%. With a sexual transmission rate of 0.0175 (corresponding to about 10% probability of transmission per coital act), the attack rate increases to 15.97%. This accounts to about 10.29% increase in the epidemic size that can be caused by a high rate of sexual transmission. We also have an unknown factor, the relative sexual transmissibility (*μ*) when recovering hosts are in the convalescent phases. For all other simulations, we assume the infectiousness will reduce by 50% in convalescent stages. However, we also vary this parameter to determine how it affects the outcomes. Just like the effects we saw for *β*, the epidemic lengths remain within 1-2 days of the mean of 157.6 days (Figure 5d). The pathogen survival also show similar characteristics as before and stays within 73-76 days (Figure 5e). The effect on the epidemic size is barely noticeable (Figure 5f), with small fluctuations.

**Figure 5.**
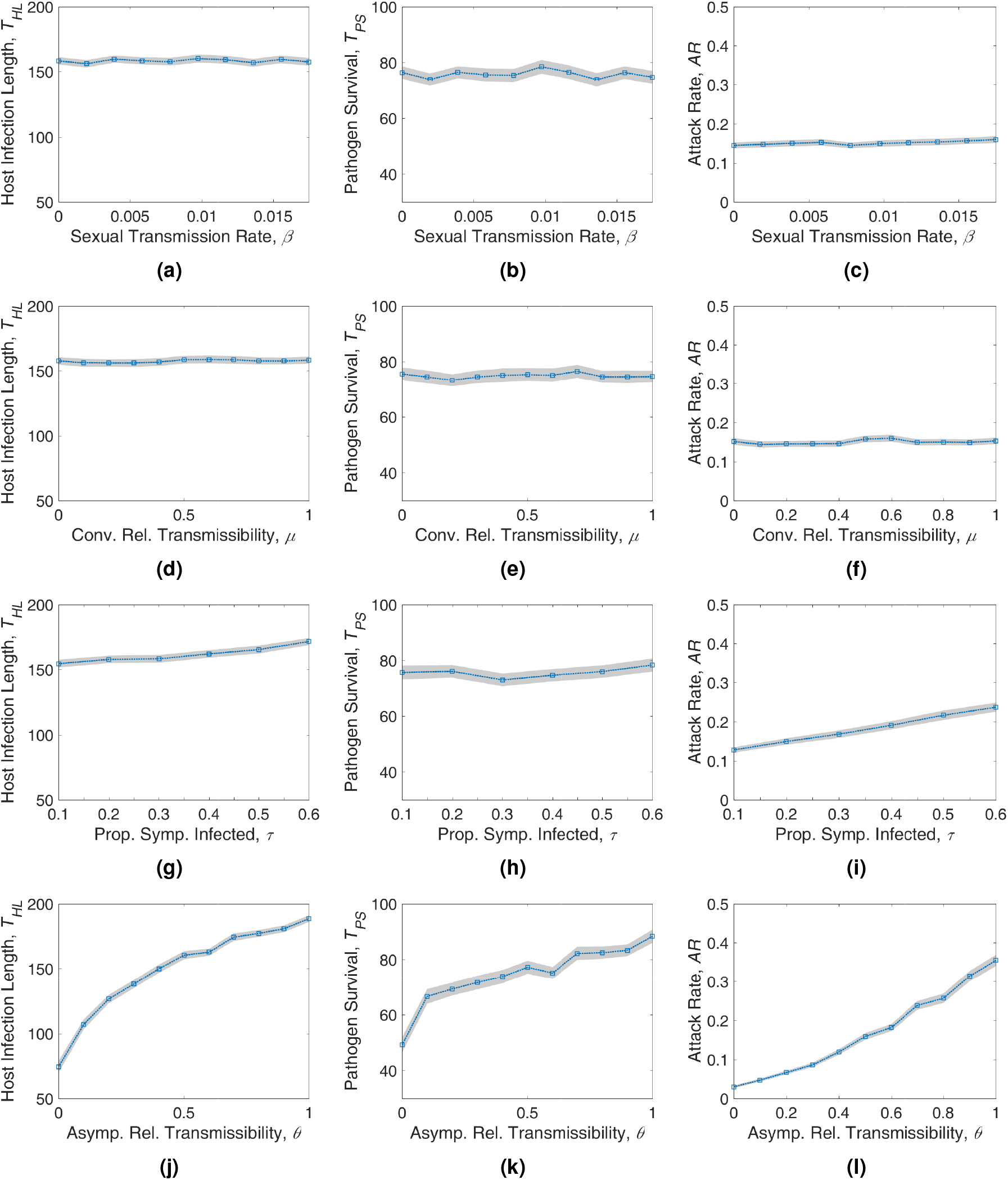
The plots depicting the results of pathogen survival analysis. The results were obtained by varying the sexual transmission rate *β* (1^st^ row), the relative transmissibility of the convalescent states *μ* (2^nd^ row), the proportion of symptomatically infected individuals *τ* (3^rd^ row), and the relative transmissibility of the asymptomatic states *θ* (4^th^ row). The plots demonstrate the host infection length *T_HL_* (1^st^ Col), the pathogen survival *T_PS_* (2^nd^ col), and the attack rate *AR* (3^rd^ col). Each data point (blue square) in the above plots is an average of 500 independent stochastic simulations. The shaded regions indicate 95% confidence intervals.

ZIKV disease outcomes are also affected by a large proportion of asymptomatic hosts. We vary the two parameters (*τ* and *θ*) that we use to model the asymptomatically infected individuals. Although in most works we found that about 20% of the cases were symptomatic, one recent work estimated that this proportion could be higher29 (27% - 50 %). To evaluate the effect of such variations, we vary the symptomatic proportion parameter, *τ* from 10% to 60%. The epidemic length increases gradually with τ starting from 155 days for 10% symptomatic to 172 days for 60% symptomatic population (Figure 5g). A 50% increase in the symptomatic proportion causes a 11% increase in the length of the outbreak. The pathogen survival is affected much less compared to epidemic length. However, we find a small fluctuating decrease followed by a gradual increase (Figure 5h). The vector free pathogen survival remains within 2-3 days of the average length of 75.6 days. The epidemic size has a clear increasing trend as shown in Figure 5i. This is expected because the symptomatically infected individuals are subject to higher transmission rates. As an example, the outbreaks will be about 45% larger if 50% of the people are symptomatic compared to the usual proportion of 20%. The relative transmissibility has significantly larger effect in the disease outcomes and all three properties increase (Figure 5 bottom row). In most other simulations, it is assumed by default that infectiousness reduces to half (θ = 0.5) when someone is asymptomatic. Compared to our default situation, if we make infectiousness of both symptomatic and asymptomatic individuals the same, we see 17.5% increase in the epidemic length (Figure 5j), 14.45% increase in the pathogen survival period (Figure 5k), and a massive 123.5% increase in the epidemic size (Figure 5l).

#### Sensitivity Analysis

A sensitivity analysis is performed in order to evaluate the model response with respect to several key parameters. For a vector borne disease such as ZIKV, the ratio of vector and host population is expected to play a prominent role in the disease outcomes.

In our previous analysis, we have also found that the proportion of asymptomatic individuals and their relative transmission rate affect the disease outcomes. To evaluate how these parameters interplay, we perform a pairwise sensitivity analysis of the three parameters: the vector-to-host ratio (*N_V_/N_H_*), the proportion of asymptomatic hosts (*τ*), and the relative transmissibility of asymptomatic hosts (*θ*). The results are depicted in Figure 6.

**Figure 6.**
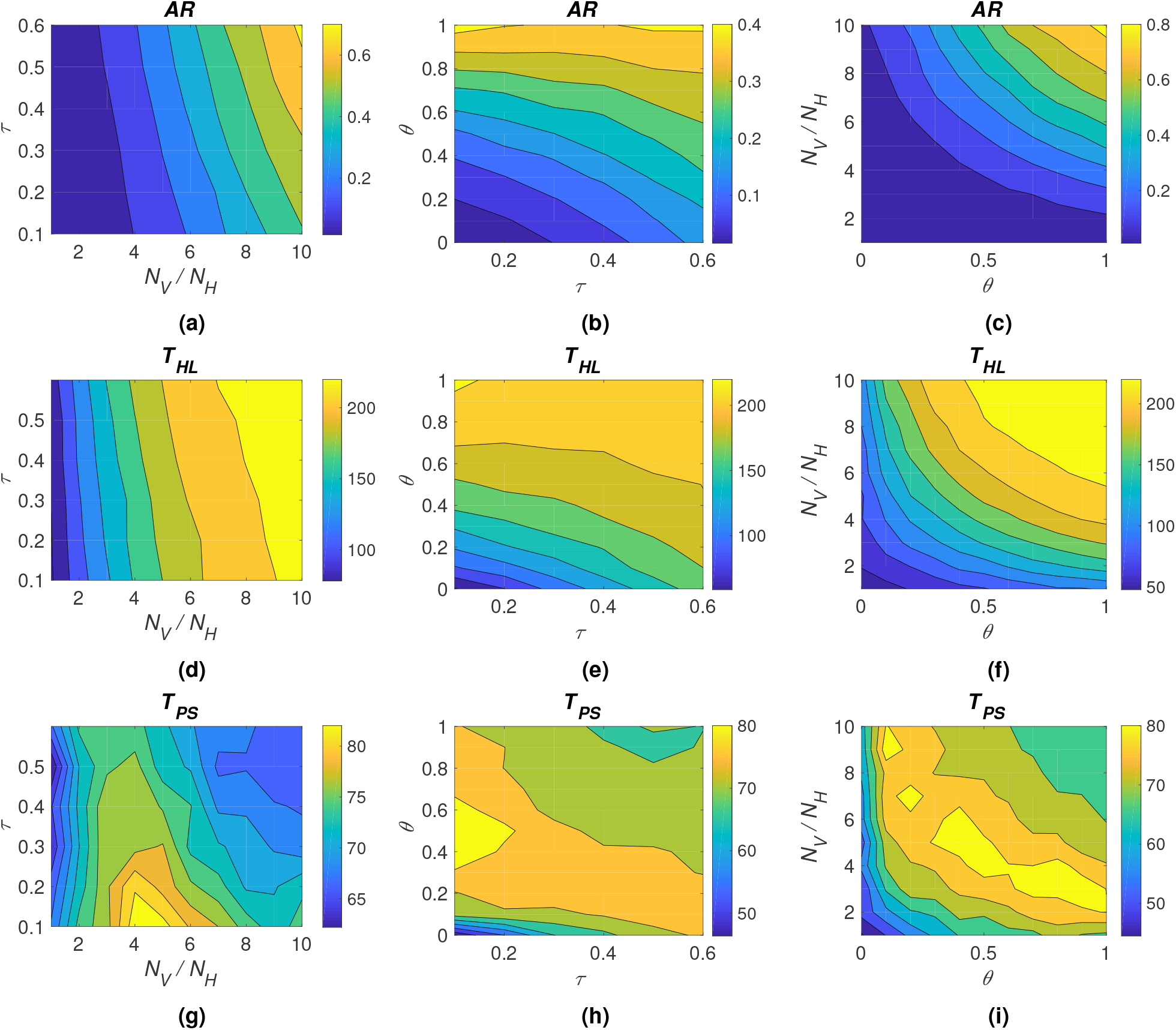
Contour plots depicting the results of the sensitivity analysis. The parameters varied in each plot are marked as axes labels. The quantity for which the contours are being displayed is mentioned on top of each plot. The contours are color mapped and a color-bar on the side of each plot indicates the range of values represented by the plot. The 1^st^ row shows the sensitivity of the epidemic attack rate (*AR*) on the parameters, the 2^nd^ row shows the sensitivity of the epidemic length (*T_HL_*) on the parameters, and the 3^rd^ row shows the sensitivity of the pathogen survival (*T_PS_*) on the parameters. The parameters that were varied here are: the vector-to-host ratio (*N_V_/N_H_*), the proportion of symptomatically infected individuals (*τ*), and the relative transmissibility of the asymptomatic states (*θ*). Each data point shown in the above plots is an average of 500 independent stochastic simulations.

The epidemic attack rate (*AR*), as expected, is highly sensitive to vector-to-host ratio, *N_V_*/*N_H_* for the entire range of asymptomatic proportion, *τ*. For the nominal value of *τ* = 0.2, doubling the *N_V_/N_H_* ratio increases epidemic size by 249.78% (Figure 6a). However, if the asymptomatic individuals have very low transmission capabilities, it can limit the effect of vectors on epidemic size as seen in Figure 6c. The fact that majority (80%) of the hosts are asymptomatic in ZIKV infections, indicate that they hold a critical role in spreading. The parameters *τ* and *θ* both demonstrate importance in determining epidemic size (Figure 6b) and among them, the effect of *θ* is more radical. Notice, the color bars on the right of the plots do not have the same scales, hence the plots can’t be compared side by side. Instead, we are looking at relative impacts of parameters in pairs.

The epidemic length (*T_HL_*) is also affected by the three parameters in question. However, we can see that, that effect of symptomatic proportion depends on the relative transmissibility. The symptomatic proportion can regulate epidemic length if relative transmissibility is low as seen in Figure 6e. If *θ* is above 0.4, the capability of *τ* is greatly reduced. The vector-to-host ratio (*N_V_*/*N_H_*) always increases the epidemic length (Figures 6d and 6f), which is also true for the relative transmissibility of asymptomatic individuals *θ* (Figures 6e and 6f).

We obtain more interesting results when we analyze the sensitivity of pathogen survival, *T_PS_.* Pathogens can survive longer for an intermediate range of host to vector ratio if the proportion of symptomatic individuals is low as shown in Figure 6g. On the other hand, despite having some large epidemics on the upper right corner of Figure 6a, we see that pathogen survival is relatively lower (as low as 67 days) (Figure 6g). In Figure 6h, we see that depending on the value of *τ*, the survival can be higher for certain intermediate values of *θ*. For example, When *τ* < 0.2 and 0.3 < *θ* < 0.7, there is a region where pathogen can survive more compared to the other situations. Survival reduces for both high and low values of asymptomatic relative transmissibility (*θ*). When comparing vector-to-host ratio along with the asymptomatic relative transmissibility, we see that a thick band or a region appears, where the pathogen survival is high (Figure 6i). There are slight fluctuations in that peak region, however, they are relatively minor to indicate any particular phenomena. If we compare this plot with the attack rate plot (Figure 6c), we see that pathogen survival is longer for intermediate values of the attack rates and shorter for the very small or very large outbreaks. Small outbreaks naturally ends sooner. Very large outbreaks also can end soon due to faster spreading dynamics and herd immunity of the recovered population.

## Discussion

In this study we have proposed an individual based interconnected network model for ZIKV that can also be used to simulate any vector-borne disease that features contact based direct transmissions. We employ heterogeneous mixing based on the host contact network which is generated based on real world data on human sexual behavior, sexual orientation, gender, and age structure. We utilize the approximations of the non-Markovian Gillespie algorithm to run stochastic simulations. In the beginning, we explore how the seasons can affect this predominantly vector-borne disease. Later, we focus on the survival of pathogen in the climates similar to Florida if an outbreak starts prior to the colder months. In this step, we examine how some of the important model parameters affect the outcomes. Finally, we perform a sensitivity analysis to evaluate our model behavior in response to changes in some key parameters.

Our seasonal analysis results indicate that outbreak size is strongly related to the environmental conditions during the pathogen introduction. The first few weeks are crucial in determining how much the pathogen would spread. After that period, environmental variations have a much weaker effect in reducing or increasing the outbreak size. If the pathogen is introduced during the peak mosquito season, there is a high probability that we will see a large outbreak. Even if climatic suitability of mosquito vectors decline rapidly after the first few weeks, the vectors manage to spread the pathogen in the host population rapidly and cause large outbreaks. This suggests that a ZIKV outbreak can spread rapidly out of control if it is not effectively contained during the initial stage. Early interventions are crucial even though it may be difficult to identify outbreaks due to large asymptomatic group of hosts. The ratio of vector to host, as expected, has shown its prominence in determining outbreak size and the length (Figure 6).

We have analyzed how sexual transmission affect the outbreaks. Our results in Figures 5c and 5f align with the conclusions drawn by some of the earlier works^4–6^ that sexual transmission is a small component in the overall force of infection, which is dominated by the vectored-transmission. However, pathogen still survives up to 75 days in the host network without the help of infected vectors. This long survival can be attributed to small amount of sexual transmission but it is mainly due to the extended infectiousness (convalescence) of the hosts as it is assumed that pathogen can survive up to a month in the semen of recovering males. The use of certain birth controls (e.g. condoms) could be effective in combating the spread of pathogen during this period. These conclusions should inspire further clinical studies in this area to test the efficacy of control measures.

A large number of asymptomatically infected individuals play an important role in outbreak dynamics. The proportion of symptomatic individuals do not affect the survival of pathogen noticeably. However, it is positively correlated to outbreak size. The relative transmissibility of asymptomatic states is one of the key factors in determining disease outcomes which can extend the vector-free pathogen survival up to 3 months. It is one of the most important parameters that needs to be properly estimated in order to obtain informative outbreak predictions.

The conclusions drawn from our results can be useful in evaluating potential endemic scenarios for Zika virus disease in temperate regions. Although the contribution of sexual transmission is small, the ability of the pathogen to survive in a human sexual contact network for extended periods can have consequences in sustaining a Zika outbreak in a region as well as spreading to other regions due to long distance travels. In addition to that, a large asymptomatic proportion could be the most important hurdle in controlling ZIKV outbreaks. Although we provide conclusions on the relative importance of key parameters, the unavailability of data on some of those warrants for further efforts towards estimations. This model should be applicable to other vector-borne disease which have the potential of being transmitted sexually. Our model is also mesoscale (medium sized population) in terms of hosts, due to the limitations imposed by computational complexities. This work can be extended in the future by the use of activity driven networks *(ADN),* vertical transmission, and larger populations. Those studies would provide more insights into the study of Zika virus epidemics.

## Methods

### Host Network Characterization

For the host population, we assume equal number of males and females. We divide the population into three age groups: Children (0-14), Adults (15-64), and Elderly (65+). Based on the World Development Indicators (WDI) data published for 2017^36^, we find that for the USA population, the three age groups are distributed as 19% Children, 66% adults, and 15% elderly people. The adults are the only ones assumed to be capable of transmitting the disease sexually, hence the remaining population are only affected by vectors. Based on the data provided in a sexual behavior study^23^, we find that sexual orientation of men consisted of approximately 97.2% heterosexuals, 2.5% homosexuals and 0.3% bisexuals. For women, the study revealed 98.9% heterosexuals, 0.9% homosexuals and 0.2% bisexuals. The same work23 also compiles a distribution of the population based on the number of sexual partners which is listed in Table 2. The data indicate that majority of the population have a single partner in a timespan of 12 months. The average number of partners for the adult population was 1.28.

**Table 2.**
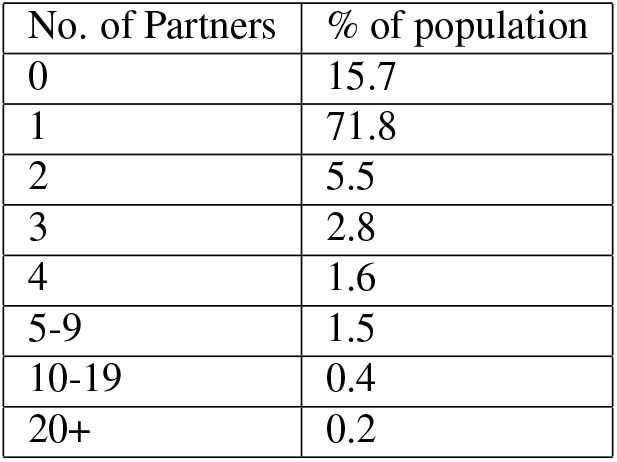
Average number of sexual partners in the last 12 months^23^

The network generator tool is designed using the configuration model^34^. It takes all the above mentioned population and sexual behavior properties as inputs, and generates a graph whose statistical properties closely match the given partner distribution of Table 2 and the distribution of sexual orientation. Studies on sexual networks also indicate that sexual contact networks have node degree distributions that follow the scale-free structure^37^. There are large variations in the number of sexual partners while a small group of highly active people form the core^37^. Core groups are important in sustaining pathogen transmission specially if the duration of infection is short^38^. To maintain these features, the network generator is designed in such a way that, high degree nodes have higher probabilities of connecting to low degree nodes. An example of network generated by our generation tool is shown in Figure 3.

### Vector Characterization

The mosquito vectors are modeled as homogeneous population and we use the classical Ross-Macdonald approach used by Keeling et al. in their book^39^. We divide the vector population into three compartments, Susceptible (*S_V_*), Exposed (*E_V_*) and Infected (*I_V_*). The transitions between the three compartments are showed in Figure 1. Table 1 describes the different parameters that were used for the model. The equations for disease dynamics in mosquito vectors are given below,

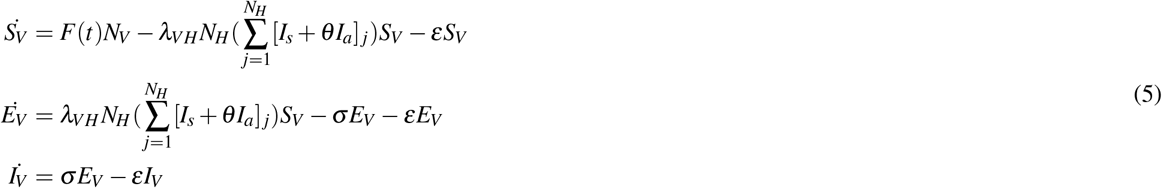

We incorporate seasonality into this model by using a time dependent mosquito recruitment rate, *F*(*t*). This rate depends on the time (day) of the year. The transmission parameters, the *λ*’s are computed from the mosquito bite rate, *r* and transmission probability, *T*. The formula are given in Table 1.

### Numerical Simulation

We use the non-Markovian Gillespie Algorithm (nMGA) to simulate the processes related to host nodes. The vector population is simulated using deterministic ordinary differential equation (ODE) solver. We combine the hosts and the vector simulations together by calling the ODE solver at the end of each event. The GEMFsim27 tool, which already supports the Gillespie algorithm was modified in order to include the vector population model and time varying parameters. We use the variables *X* and *Y* to describe the host and vector populations in different states/compartments respectively. Here, *X* contains information about the state of each host node and *Y* contains the population count of each vector compartment. *X*_0_ and *Y*_0_ are the initial states/compartments at the start of the simulation. 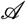 is the host network representation. The term *PARAM* is used to denote the set of model parameters listed in Table 1. *T_H_* and *T_V_* in the output are the host and vector indexing arrays that contain information about time where the data points (*X* and *Y*) were generated. We denote *x_n_* ∈ *X* as the state of node *n. A_n_*(*x_n_* → *j*) is the transition rate of node *n* from its current state *x_n_* to state *j*. We use *M* to denote the total number of host states (= 4 in our case). For a node *_n_*, Λ_*n*_ denotes the sum of transition rates from its current state to all other possible states. The *VECTSOLVER* is an ordinary differential equation (ODE) solver that takes as input the current situation of the vector population *Y* and solves them from the current instance to the time *δt*. The ODE equations that are being solved are given in Equation 5. This solver is invoked once after each event (with the updated parameters) and the time indexing vectors are updated with respect to the global time *t*. The simplified pseudo-code of the modified GEMFsim is given below,

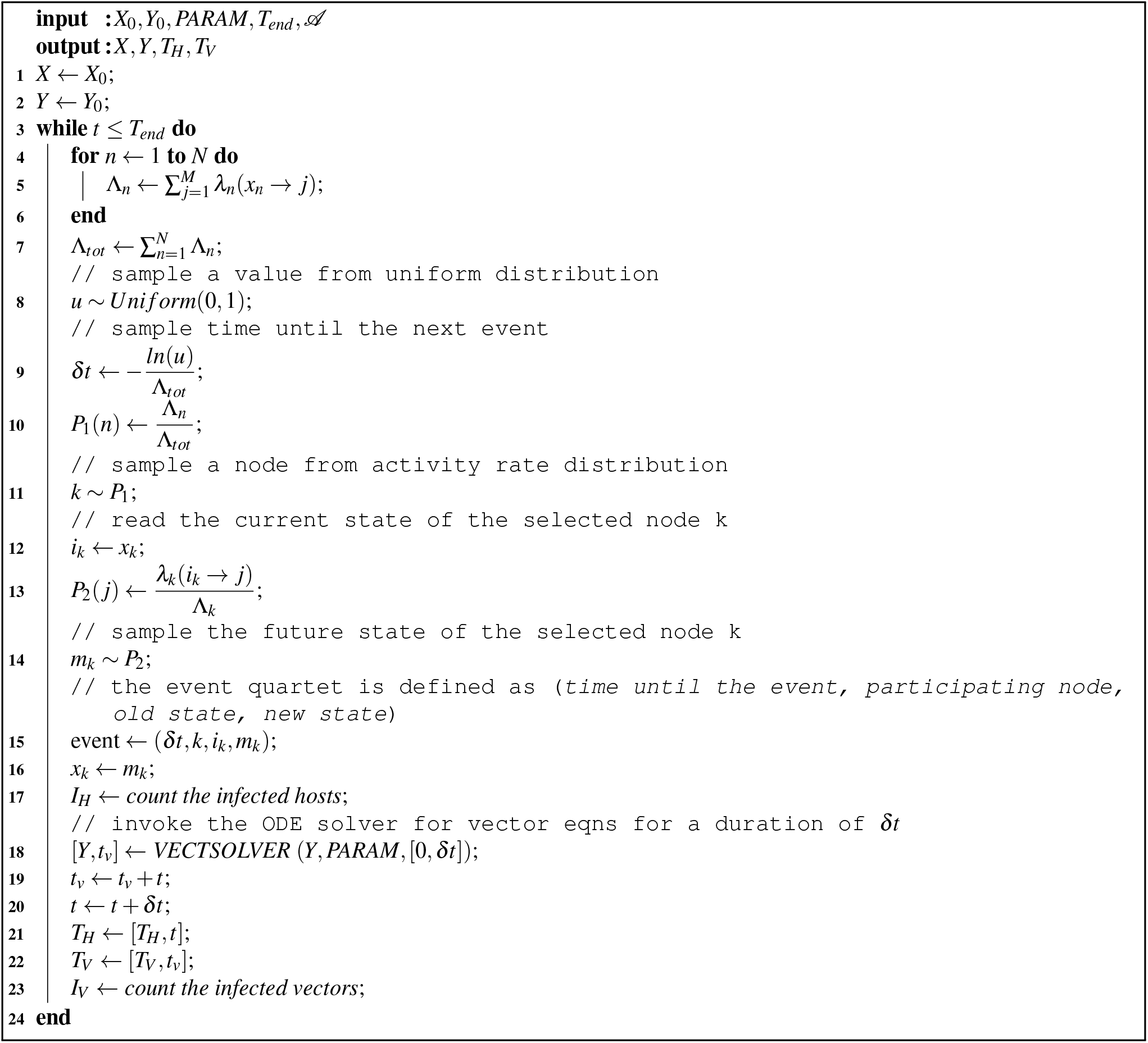

As the simulation is event-based and parameters are updated following each event, the inter-event times should be small to keep the variable parameters up to date in both populations. Fortunately, due to the nature of the Gillespie algorithm, it is indeed the case when a pathogen is present in the host population. We have demonstrated this fact in Figure 7 that ensures parameter accuracy. The inter-event times are also exponentially distributed which is shown in Figure 7b.

**Figure 7.**
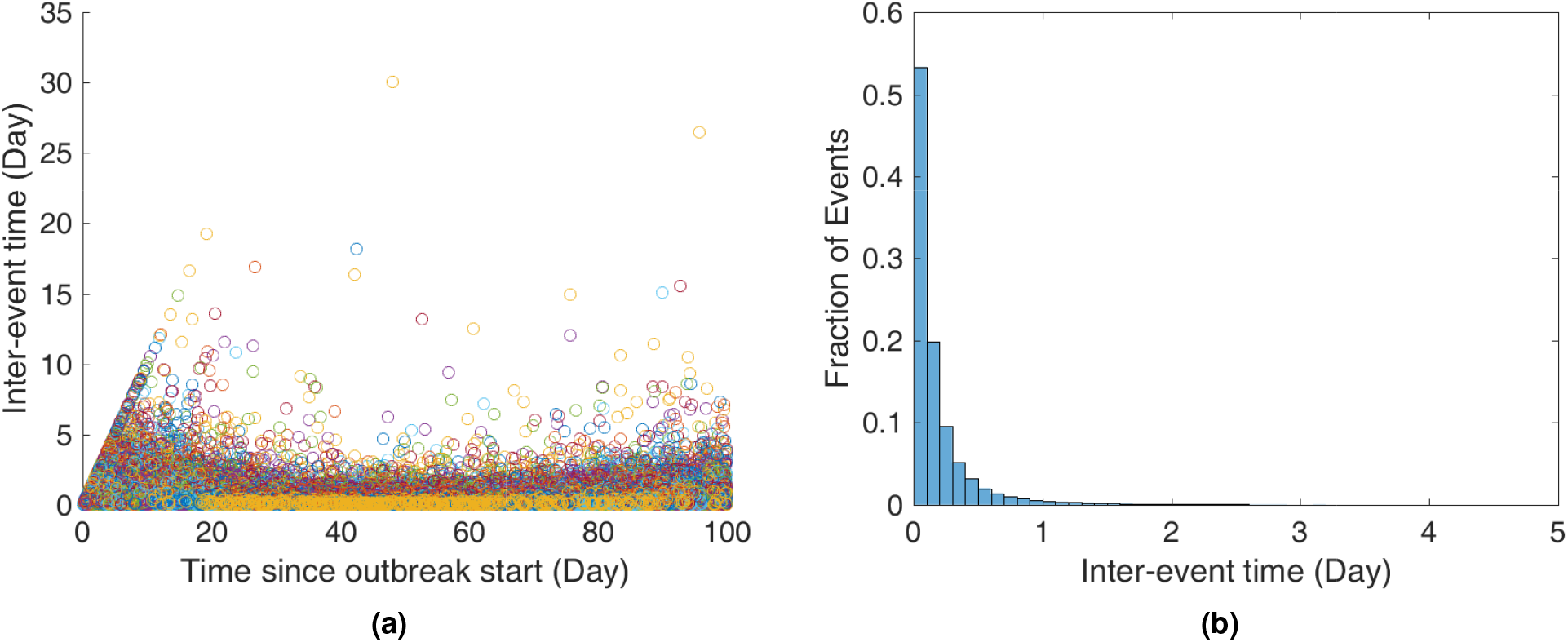
Inter-event times in different stages for all of the 500 simulations during the first 100 days (left) and their distribution (truncated inter-event times > 5) (right). For this case, the average inter-event time is 0.2074 day (95% CI [0.2056 to 0.2092]). About 96.87% of the events that occurred had intervals shorter than a day. There are a few outliers though, which are mostly due to slowing down of the events at the end of the epidemics (when pathogen is low in the host population or have been wiped out).

### Non-Markovian Gillespie Algorithm

The nMGA (Non-Markovian Gillespie Algorithm) was proposed by Boguna et al^26^. The following derivation was also described in the work of Masuda et al.^40^.

We first consider *N* renewal processes running in parallel. Let *t_i_* be The time elapsed since the last event of the ith process. We denote *ψ_i_*(*τ*) as the probability density function of inter-event times for the *i*th process. The survival function of the ith process (i.e., the probability that the inter-event time is larger than *t_i_*) is,

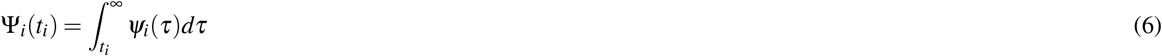

Now, the probability that no process generates an event for time Δ*t* is,

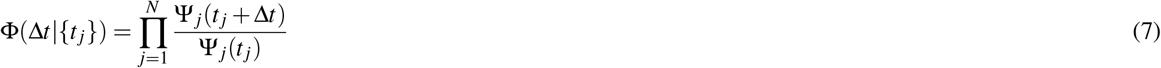

To determine the time until the next event, Δ*t*, we take a sample *u* from uniform distribution over [0, 1] and solve Φ(Δ*t* |{*t_j_*}) = *u*. This step is computationally expensive when *N* is large. To improve performance, we approximate this step as proposed by Boguna et al^26^. This approximation is exact as *N* → ∞. When Δ*t* is small (*N* is large), Equation 7 becomes^40^,

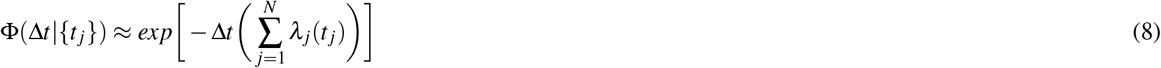

Now, the instantaneous (hazard) rate of the *i*th process, which is generally assumed to be a function of time since the last event is determined by,

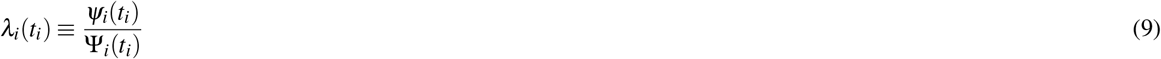

With the above equations in hand, we can run the Non-Markovian Gillespie algorithm as follows,

1. Initialize *t_j_* for all (1 ≤ *j* ≤ *N*).
2. Determine the time until next event from,

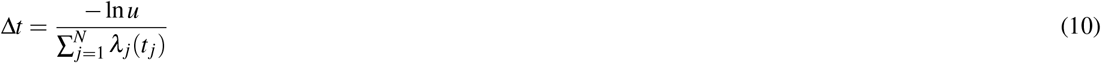
3. Select the process *i* that has generated the event with probability,

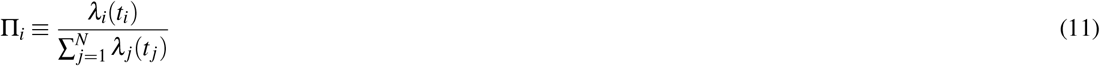
4. Update the time since the last event, *t_j_* = *t_j_* + Δ*t* for all *j* ≠ *i*. Set *t_i_* = 0.
5. Repeat steps 2-4.

The original Gillespie Algorithm can be recovered from this nMGA using *λ_i_*(*t_i_*) = *λ_i_*. This is the case for all the parameters that are constant.

## Acknowledgements

The authors gratefully acknowledge the financial support provided by the National Science Foundation under Grant Award CIF-1423411, United States Department of Agriculture under the Grant No. 2015-67013-23818 and 3020- 32000-008-04-S.

## Author contributions statement

T.F., C.S., and L.C. formulated the model; T.F., L.C., and D.M. analyzed the data; T.F. and C.S. developed the methodology; T.F. developed the simulation tool; T.F., D.M., and L.C. analyzed the results and contributed to the discussions; T.F., L.C., D.M., and C.S. contributed to the writing. All authors reviewed the manuscript.

## Additional information

### Competing interests

The authors declare that they have no competing interests.

